# Improving reliability in clinical neuroimaging: a study in transgender persons

**DOI:** 10.1101/861864

**Authors:** Behzad Sorouri Khorashad, Behnaz Khazai, Ali Talaei, Freya Acar, Anna R. Hudson, Nahid Borji, Hedieh Saberi, Behzad Aminzadeh, Sven C. Mueller

## Abstract

Although the neuroanatomy of transgender persons is slowly being charted, findings are presently discrepant. One important factor is the issue of power and low signal-to-noise (SNR) ratio in neuroimaging studies of rare study populations including endocrine or neurological patient groups. The present study assessed whether the reliability of findings across structural anatomical measures including thickness, volume, and surface area could be increased by using two back-to-back within session structural MRI scans in 40 transgender men (TM), 40 transgender women (TW), 30 cisgender men (CM), and 30 cisgender women (CW). Overall, findings in transgender persons were more consistent with at-birth assigned sex in brain volume and surface area while no group differences emerged for cortical thickness. Repeated measures analysis also indicated that having a second scan increased SNR in all ROIs, most notably bilateral frontal poles, accumbens nuclei and putamina. Furthermore, additional significant group differences emerged in cortical surface area when age and ICV were used as covariates. The results suggest that a simple time and cost effective measure to improve signal to noise ratio in rare clinical populations with low prevalence rates is a second anatomical scan when structural MRI is of interest.

## Introduction

Mounting research efforts over the past two decades have been trying to determine whether transgender persons bear more neuroanatomical resemblance to their gender identity vs. their at-birth assigned sex [1–3]. Yet, neuroanatomical findings in transgender persons remain highly discrepant possibly due to a variety of factors including sexual orientation [4,5], genetics [6], hormonal factors [7] as well as small size of the cohorts [8]. In addition, cultural factors have remained unexplored to date as the majority of transgender neuroimaging has been conducted in Western populations [2].

In terms of neuroanatomical differences between the sexes, large-scale studies and meta-analyses have tried to identify typical anatomical patterns for cisgender men (male at-birth assigned sex and male gender identity) and cisgender women (female at-birth assigned sex and female gender identity)[9,10]. These findings are regionally specific but curiously, also partly discrepant. For example, Ruigrok et al. [10] reported that, on average, males have larger grey matter volume (GMV) in the putamen and the cerebellum, whereas females have, on average, larger volume of the right frontal pole and parietal cortex. Interestingly, while the meta-analysis of Ruigrok et al. [10] from several different cultures indicated larger thalamic volume in women relative to men, the single population study (with a larger sample size) of British participants by Ritchie et al. [9] documented larger thalamic volume for men relative to women. Highly regionally-specific findings are further exemplified by a recent analysis which showed it to be rare that brains consistently belong to either extreme ends of the maleness-femaleness spectrum are rare and that they rather are “mosaics” of male and female features [11]. This mosaic model could be further supported by the findings of Lombardo et al. [12] who showed an association between fetal testosterone level- as measured in utero through amniocentesis- and GMV in bilateral somatosensory, motor, and premotor cortices in a cohort aged 8-11 years old. Moreover, both Lombardo et al. [12] -using a human sample-and [13]-using an animal sample-showed that inferior parietal lobule is sensitive to androgens.

With regards to the “maleness-femaleness” spectrum, Guillamon et al. [1] postulated that trans people demonstrate different extents of phenotypically male and female brain patterns, with trans women displaying some demasculinized patterns and trans men some defeminized patterns. This yields their own unique brain patterns and possibly shifting trans men’s brains towards those of cis men and trans women’s brains towards those of cis women. However, findings have been inconsistent. Some studies show female-typical brain patterns for trans women in hypothalamus [14], left pre- and postcentral gyri [15], or nucleus accumbens [16], while others find brain patterns consistent with the sex-assigned at birth in putamen, precentral gyrus [17] and frontal lobe [18]. Such inconsistencies are found among different anatomical measures including GMV and cortical thickness [16,19–21], as well as cortical surface area [16,20]. Ongoing extensive research is required to achieve consistency and conclusive results.

Yet, as noted above, an important caveat to bear in mind is the typical small sample sizes used in neuroimaging studies of transgender persons which imposes great limitation to reach definitive conclusions regarding anatomical patterns. While this factor is inherent to cohorts with small prevalence, prior work in healthy and neurological cohorts has suggested that using a second MRI scan might improve power, Signal-to-Noise Ratio (SNR), and reliability [22–24]. In a large cohort of healthy elderly participants (64+ years), Liem et al. [23] examined reliability within sessions of two anatomical scans taken 30 minutes apart and concluded that variance due to measurement error can be decreased and SNR increased by simply scanning additional participants. The Alzheimer’s Disease Neuroimaging Initiative (ADNI,[25] has also demonstrated the utility of additional scans in a neurological cohort. However, to our knowledge the effect of direct back-to-back scans has not been examined explicitly.

*Therefore, the study had 2 main objectives. The first objective was to replicate prior neuroimaging studies in transgender persons in a Middle Eastern (Iranian) population considering the lack of comparative data in Non-Western cultures. Building on the prior achievements in healthy [23] and neurological cohorts [25], the second objective was to increase SNR and confidence in findings by collecting an additional second anatomical scan.* Therefore, we focused on 10 *a priori* regions of interest that have been suggested to differ between cisgender males and females and also 2) implicated in prior reports of transgender neuroimaging [16,17,26–28]. These ten regions of interest (ROI) were the fusiform gyri, inferior parietal gyri, pre- and postcentral gyri, frontal poles, thalami, caudates, putamina and accumbens nuclei as well as the cerebellum.

## Results

### Demographics

There were no significant differences in age between TM (Mean age = 24.37 years; SD = 5.35 years), TW (Mean = 24.87; SD = 6.2), CM (Mean = 26.03; SD = 5.25) and CW (Mean = 25.8; SD = 3.83) (Table 1). While there was no significant difference between CM and CW as well as TM and TW in their socioeconomic status (SES), cisgender participants had significantly higher SES compared to transgender participants (χ(2) = 9.86, p =.007). A similar pattern emerged for the educational status: cisgender and transgender participants differed from one another (χ(3) = 25.41, p < .001), but not within groups. More importantly, with regards to gender identity and sexual orientation, TM reported to have significantly more masculine gender identity compared to CM (t (68) = 2.27, p = .033) but they did not differ in sexual orientation (both were predominantly attracted to women), while TW and CW did not differ in either sexual orientation (both were predominantly attracted to males) or gender identity (both held feminine gender identity). Only two people among the whole sample scored 7 on the sexual orientation scale - one cis man and one cis woman - and no one scored in the 4-6 range, which suggests a binary distribution of sexual orientation in this sample.

**Table 1.**
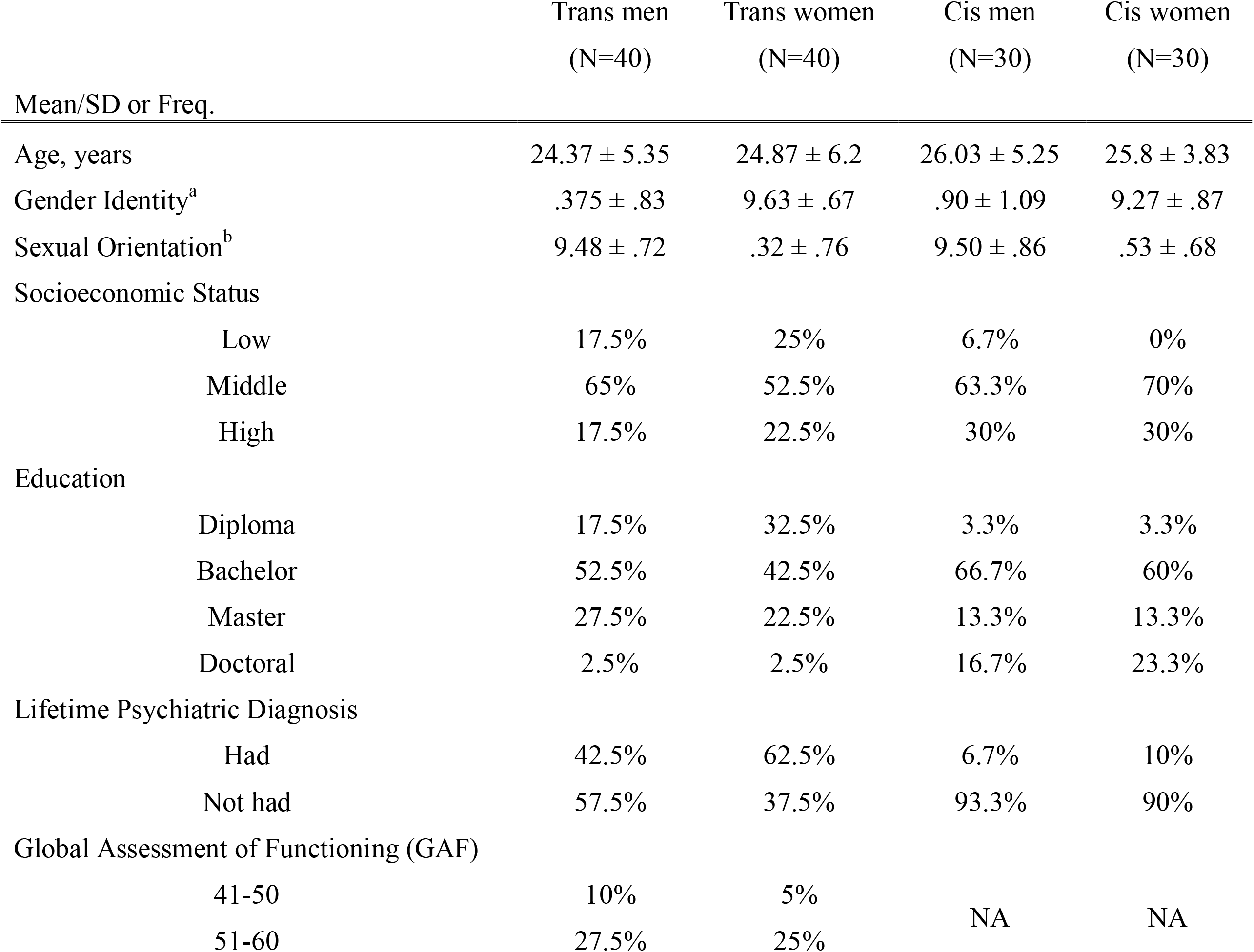
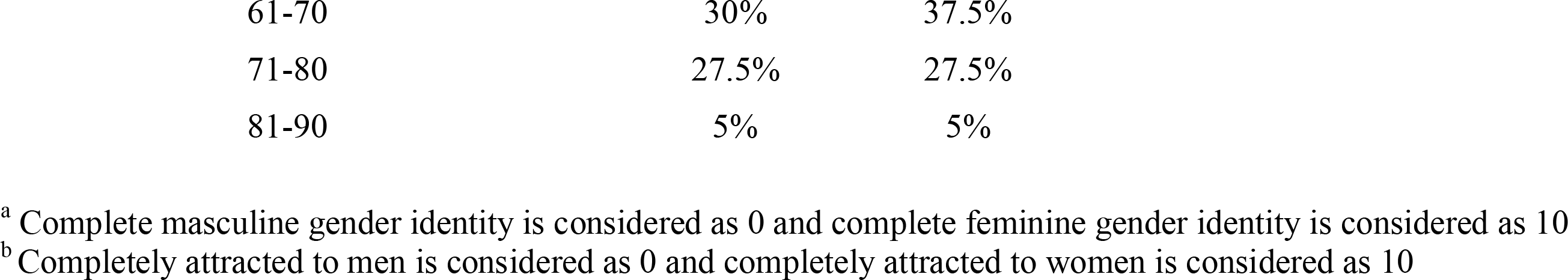
Demographic information of TM, transmen; TW, transwomen; CM, cisgender men; CW, cisgender women

|  | Trans men<br>(N=40) | Trans women<br>(N=40) | Cis men<br>(N=30) | Cis women<br>(N=30) |
| --- | --- | --- | --- | --- |
| Mean/SD or Freq. |  |  |  |  |
| Age, years | 24.37 ± 5.35 | 24.87 ± 6.2 | 26.03 ± 5.25 | 25.8 ± 3.83 |
| Gender Identity<sup>a</sup> | .375 ± .83 | 9.63 ± .67 | .90 ± 1.09 | 9.27 ± .87 |
| Sexual Orientation<sup>b</sup> | 9.48 ± .72 | .32 ± .76 | 9.50 ± .86 | .53 ± .68 |
| Socioeconomic Status |  |  |  |  |
| Low | 17.5% | 25% | 6.7% | 0% |
| Middle | 65% | 52.5% | 63.3% | 70% |
| High | 17.5% | 22.5% | 30% | 30% |
| Education |  |  |  |  |
| Diploma | 17.5% | 32.5% | 3.3% | 3.3% |
| Bachelor | 52.5% | 42.5% | 66.7% | 60% |
| Master | 27.5% | 22.5% | 13.3% | 13.3% |
| Doctoral | 2.5% | 2.5% | 16.7% | 23.3% |
| Lifetime Psychiatric Diagnosis |  |  |  |  |
| Had | 42.5% | 62.5% | 6.7% | 10% |
| Not had | 57.5% | 37.5% | 93.3% | 90% |
| Global Assessment of Functioning (GAF) |  |  |  |  |
| 41-50 | 10% | 5% | NA | NA |
| 51-60 | 27.5% | 25% |  |  |

|  |  |  |
| --- | --- | --- |
| 61-70 | 30% | 37.5% |
| 71-80 | 27.5% | 27.5% |
| 81-90 | 5% | 5% |

<sup>a</sup> Complete masculine gender identity is considered as 0 and complete feminine gender identity is considered as 10
<sup>b</sup> Completely attracted to men is considered as 0 and completely attracted to women is considered as 10

For mental health status, 91.7% among cisgender participants (n = 60) reported to have no current or past psychiatric condition. Five individuals reported that they had been diagnosed with depression (n = 3), general anxiety disorder (n = 1) and obsessive-compulsive disorder (n = 1). By comparison, 40% of transgender individuals had been diagnosed with at least one psychiatric condition in their axis I. Most common were Trauma- and Stressor- related disorders (20%), Depressive Disorder (11.2%), or Obsessive-Compulsive and related disorders (8.8%). There was no difference between TM and TW in their axis I psychiatric diagnoses. In the axis II, 83.8% had no diagnosis at all, 6.2% had borderline personality disorder, 3.8% had antisocial personality disorder, 3.8% had narcissistic personality disorder, one had histrionic personality disorder and one had personality disorder NOS (Not Otherwise Specified). No significant difference was found between TM and TW in their axis II diagnoses. Scores for the global assessment of functioning (GAF) among transgender participants as determined through SCID-5 is presented in Table 1.

### Anatomical differences between transgender and cisgender persons

#### GMV

For GMV, all ROIs showed significant effects of group except for the bilateral Accumbens nuclei (Table 2). Follow-up post-hoc tests are provided in Table 3. CM had larger volumes than CW in bilateral fusiform gyri, bilateral cerebella, bilateral caudates, left postcentral gyrus, right precentral gyrus, left thalamus, and right putamen. TW did not differ in any ROI from CM, who shared their sex-assigned at birth (both assigned as male), while compared to CW who shared their gender identity, they had larger left frontal pole, bilateral cerebellar cortices, right putamen and right precentral gyrus. In general, 8 out of 10 ROIs were significantly smaller in TM compared to CM, who shared their gender identity but had a different at-birth assigned sex. These regions consisted of bilateral fusiform gyri, bilateral thalami, bilateral inferior parietal gyri, bilateral pre- and postcentral gyri, bilateral cerebella, right caudate and right frontal pole. However, when compared to CW, TM who held a different gender identity but had the same at-birth assigned sex, had significantly smaller volumes in right fusiform gyrus (at trend-level in the left) and right thalamus (at trend-level in the left). Thus, they showed difference from both groups that either shared their at-birth assigned sex or their gender identity. Bilateral fusiform gyri and right thalamus findings remained significant when controlled for age and ICV (supplementary Tables 1, 2). When TW were compared directly to TM (neither shared gender identity nor at-birth assigned sex), TW had larger volumes in bilateral fusiform gyri, bilateral inferior parietal gyri, left frontal pole, right postcentral gyrus, right precentral gyrus, bilateral cerebella, bilateral thalami and right putamen; roughly consistent with the general sex effects in cisgender comparisons (Table 3).

**Table 2.** Results for group comparison for grey matter volume (GMV) and cortical surface area (CSA) without covariates (p<.05, FDR-corrected)

| Main effect of group for GMV |  |  |  | Main effect of group for CSA |  |  |  |
| --- | --- | --- | --- | --- | --- | --- | --- |
| Region | Scan 1 | Scan 2 | RM-model | Region | Scan 1 | Scan 2 | RM-model |
| L fusiform g. | **<.001** | **<.001** | **<.001** | L fusiform g. | **.002** | **.010** | **.004** |
| L inf. parietal g. | **.001** | **<.001** | **<.001** | R fusiform g. | **.016** | **.014** | **.014** |
| L postcentral g. | **.002** | **.001** | **.001** | L inf. parietal g. | **.002** | **.001** | **.001** |
| L precentral g. | **.032** | **.007** | **.012** | R inf. parietal g. | **.002** | **.004** | **.003** |
| L frontal pole | **.006** | **.045** | **.008** | L postcentral g. | **.021** | **.019** | **.019** |
| R fusiform g. | **<.001** | **<.001** | **<.001** | R postcentral g. | **.044** | .167 | .082 |
| R inf. parietal g. | **.001** | **<.001** | **<.001** | L precentral g. | **.014** | **.010** | **.012** |
| R postcentral g. | **.014** | **.011** | **.010** | R precentral g. | **.002** | **.004** | **.002** |
| R precentral g. | **.002** | **.001** | **.001** | L frontal pole | .323 | .909 | .612 |
| R frontal pole | **.001** | **.009** | **.002** | R frontal pole | **<.001** | **.004** | **.001** |
| L cerebellum | **<.001** | **<.001** | **<.001** |  |  |  |  |
| R cerebellum | **<.001** | **<.001** | **<.001** |  |  |  |  |
| L thalamus | **<.001** | **<.001** | **<.001** |  |  |  |  |
| L caudate | **.002** | **.005** | **.003** |  |  |  |  |
| L putamen | **.006** | **.013** | **.006** |  |  |  |  |
| L nucleus accumbens | .453 | .249 | .542 |  |  |  |  |
| R thalamus | **<.001** | **<.001** | **<.001** |  |  |  |  |
| R caudate | **.002** | **.002** | **.002** |  |  |  |  |
| R putamen | **<.001** | **<.001** | **<.001** |  |  |  |  |
| R nucleus accumbens | .119 | .169 | .121 |  |  |  |  |

*L: Left, R: Right; inf. =inferior; cereb. = cerebellar; g. = gyrus; RM =repeated-measures; significant p values are in bold font*

**Table 3.**
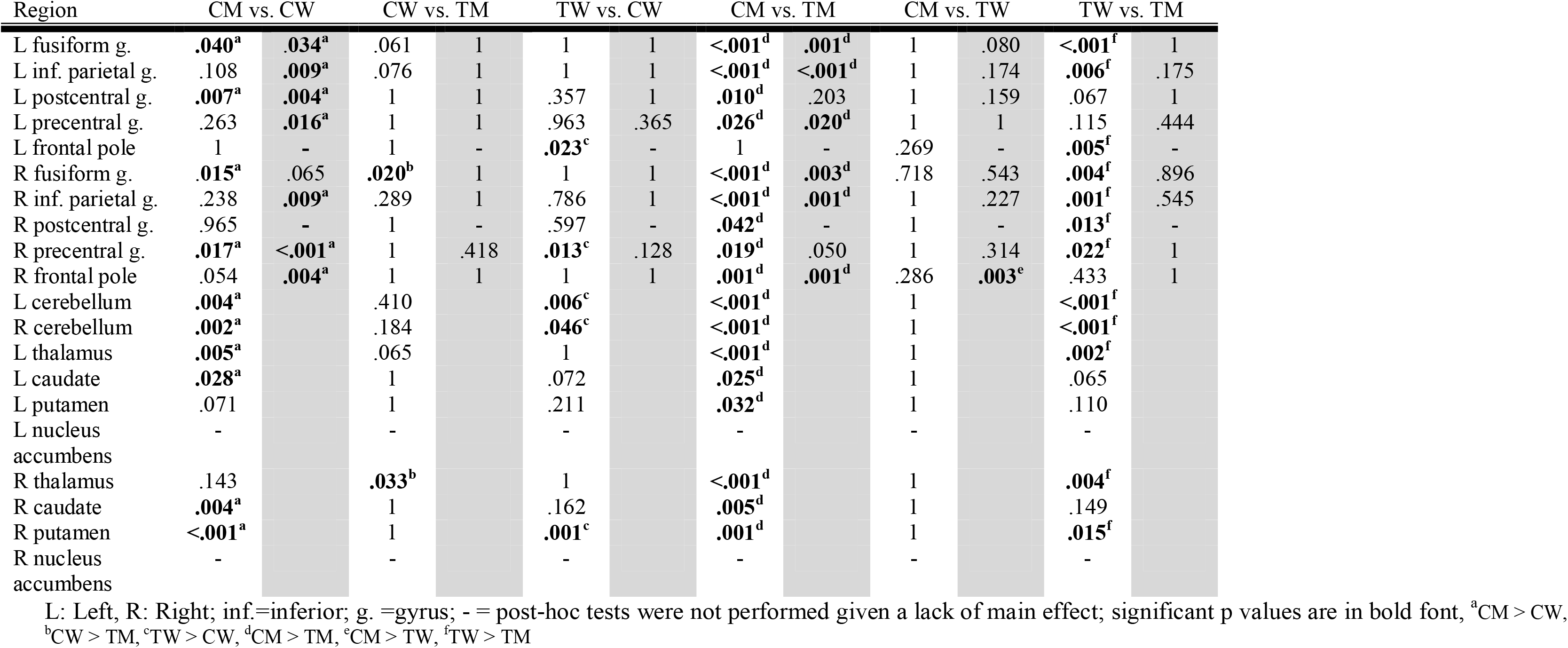
Two group comparisons for gray matter volume (GMV, in left columns, white background) and cortical surface area (CSA, in right columns, grey background) without covariates. All comparisons p<.05 (Bonferroni corrected).

#### CSA

CSA differed among the groups for all ROIs except for the left frontal pole and right postcentral gyrus (Table 2). Post-hoc analyses revealed significant differences between CM and CW, specifically, CSA was larger in CM compared to CW for all regions except the right postcentral gyrus and left frontal pole. When compared to CM, TM who had same gender identity but different at birth assigned sex, had significantly smaller surface area in bilateral fusiform gyri, bilateral inferior parietal gyri, left precentral gyrus, and right frontal pole. Right frontal CSA was also smaller in TW relative to CM but not different from CW or TM (Table 3).

### Comparing neuroanatomical measures for the two scans individually vs. a repeated measures model

The secondary objective of this study was to assess whether adding a second anatomical scan would improve reliability of our data. Primary group comparisons showed statistically significant differences in most *a priori* ROIs for GMV and CSA (Table 2). Although the repeated–measures model for main effects of group for our *a priori* ROIs did not show a substantial difference in *p*-values relative to when scans were analyzed individually (Table 2), corresponding correlations coefficients for these regions were different. Specifically, the analysis revealed a potential gain in SNR for most regions as correlation coefficients between the two MRI scans ranged between 0.705 to 0.996 (Figure 2). Interestingly, the ROIs with the lowest correlation coefficients were the bilateral frontal poles for GMV and CSA, and bilateral accumbens nuclei, left putamen and left thalamus for subcortical volume. Moreover, although adding age and ICV as covariates did not impact the main effects (supplementary Table 3), when applied in the repeated measures analysis it *did* yield more statistically significant CSA differences in all regions except for the left frontal pole and more statistically significant GMV differences for the left precentral gyrus, left frontal pole, and left putamen (Supplementary Table 1).

## Discussion

Given the current lack of comparative data [2], the main goal of this study was to assess and replicate prior findings in Western samples of transgender persons in a Non-Western (Iranian) group of hormonally naïve trans persons. Moreover, as prior inconsistencies might have suffered from small sample sizes inherent to this cohort of low prevalence, a secondary goal was to assess the potential improvement of reliability in clinical MRI data by taking two consecutive MRI scans to mitigate for low sample sizes. With regards to the first goal, strong main effects of group were present for all ROIs except for bilateral accumbens nuclei for GMV and left frontal pole for CSA. Results were generally consistent with sex-assigned at birth being the main influencer, compared to the gender identity. However, a group-specific effect emerged in the fusiform and thalamus showing lower volume for trans men relative to cis women. With regards to the second hypothesis, as expected, correlations for some regions were suboptimal, thus supporting an increase in SNR and reliability by the use of a second scan.

Endocrinological data from the ENIGI project have demonstrated the utility and necessity of providing cross-cultural data in transgender persons for the sake of generalizability. These prior data [29] have shown that sociodemographic characteristics such as the education level, age, or the amount of TM or TW presenting to the clinic may be country-specific. As with prior works from USA [26] and Sweden [17,20], the present findings across most of our *a priori* ROIs, suggest a closer neuroanatomical resemblance of transgender persons to the group they share their at-birth assigned sex with, specifically in TW. For TM, the finding was a little different, showing reduced GMV in fusiform and thalamic volume relative to CW and TM suggesting a unique profile that is different from those with shared at-birth assigned sex or gender identity. Importantly, this fusiform gyrus finding in an independent and non-Western sample has also been previously shown in a Belgian sample of hormone-treated trans persons [16], highlighting it as a cross-cultural finding in transgender individuals. Moreover, this complex pattern indicates a differential neuroanatomical profile of transgender persons relative to each other and is consistent with the hypothesis of Guillamon et al. [1]. According to their hypothesis, cortical development that has already been shown to involve thinning [21,30], is slowed/stopped in CM compared to CW, TW and TM. This thinning process affects different cortical regions and with different timings in CW, TW and TM, which leads to characteristic cortical phenotype in each group. These developmental trajectories lead to a complex mixture of masculine, feminine, and demasculinized traits in TW and masculine, feminine, and defeminized traits in TM. Therefore, this mixture may differentially affect TM and TW thus presenting with unique phenotypes. To what extent this unique fusiform gyrus and thalamus findings fits within this hypothesis remains to be investigated.

Sexual orientation may also contribute to the literature discrepancies. Guillamon et al. [1] and Savic and colleagues [4] hypothesized that sexual orientation may significantly contribute to the discrepant findings as it was not previously controlled for (e.g., [16,26] or, instead, the sample sizes were very small [15]. One strength of the present study was that both transgender and cisgender participants strongly identified with their respective gender identities and were only attracted to one other sex therefore excluding a potential confound of sexual orientation and further reducing measurement variance. Luders et al.[19] had shown thicker cortices in TW compared to CM, in an American sample of transgender persons with mixed sexual orientations, in various areas including frontal and parietal cortices with no findings in the other direction. In the present study of homogenous participants (considering their sexual orientation), we could not replicate these findings. In a better-controlled Spanish sample, Zubiaurre-Elorza and colleagues [21] reported increased CTh in TW who were predominantly attracted to men compared to CM, but this time in orbitofrontal regions, insula, and occipital cortex, with all findings being right-lateralized. Specifically probing the notion of sexual orientation in a Swedish sample, Manzouri and Savic [4] compared a heterogeneous sample (regarding sexual orientation) of TW and TM with CW who were mostly attracted to men and CM who were mostly attracted to women and found patterns consistent with shared gender identity in left-lateralized parietal and temporal cortices in TW and TM, respectively. When they reran the analyses, this time including cis women who were attracted to women and cis men who were attracted to men using Kinsey sexual orientation scores as a covariate, both the partly “female pattern” in TW and partly “male pattern” in TM disappeared. However, these findings only emerged in cortical thickness but not in volumetric measures (with surface area not being reported). In the present study, possibly because of the homogenous sample distribution, covarying for sexual orientation did not change any of the measurements of interest. Therefore, it remains to be seen whether sexual orientation or brain measurement type (or a combination of the two) is a major influence on the findings. Interestingly, an additional contributor to divergent findings might be that available studies differ in using absolute vs. relative values, and whether or not they correct for total brain volume. A case in point, sex effects between CM and CW disappeared when correcting for ICV. This may also explain some of the discrepancy between meta-analytic findings. Whereas Ruigrok et al. [10] combined (N=13) studies that either covaried for ICV (N=4), TBV (N=2), or GMV (N=7), with those that did not covary for any of these (N=3)(with 9/16 covarying for age), Ritchie et al. [9] provided both raw data measurements but computed statistics after only correcting for age. Future studies should always present both corrected and uncorrected data to facilitate future comparison.

The second goal of the present study was more of technical nature to examine how discrepancy in findings can be further reduced. Prior efforts in healthy (elderly) populations [23] and neurological cohorts such as Alzheimer’s Disease [25] have proposed that collecting additional MRI scans may improve power and reliability and increase SNR. Building on these prior accomplishments, this study aimed at examining the extent to which the literatures discrepancies in this small cohorts might have been due to the low power by collecting a second anatomical scan. Within this context, prior work in gender identity has documented that the right putamen consistently differs in volume between transgender and cisgender persons, albeit in different directions [16,17,26]. Consistent with our hypotheses and prior work, correlation coefficients were high suggesting good reliability in the FreeSurfer pipeline. However, although correlation coefficients in many regions were .90, none had a perfect correlation (except the cerebellum). Strikingly, subcortical structures such as putamen, nucleus accumbens and thalamus, as well as regions close to the edge of the brain (the frontal poles) exhibited relatively low correlations, suggesting that structural studies with a focus on these brain regions may benefit from adding a second scan to increase reliability. On the one hand, these finidings point to the need for increased sample sizes or additional scans, and on the other hand, for refinement of presently available toolboxes such as FreeSurfer to improve algorithms of parsing anatomical regions in order to increase the reliability in these areas. The present findings may be of particular interest for researchers aiming to pinpoint the underlying neural architecture of basic brain processes including reward processing [31], motor processing [32], theory of mind [33], or nociception and pain [34].

Some potential limitations of the current study merit notion. Firstly, some have expressed doubts on reliability of self-reports on gender identity or sexual orientation in Iranian samples because of the socio-political context and the Islamic republic jurisdiction. For example, as shown by Khorashad et al. [35,36], both transgender and intersex participants scored higher on the Ambivalent Sexism Inventory (ASI) compared to cisgender participants. Based on the Gender Self-Socialization Model (GSSM) by Tobin [37], Khorashad et al. [35] proposed that increased scores of ASI among individuals whose gender identity is incongruent with all or some of their physical features could be an attempt to attain gender typicality. The same concept might be at play, leading them towards exaggerating their self-reported stance on gender identity and sexual orientation spectrums. Such factors constituted another strong rationale for following a very strict protocol in this university gender clinic to confirm gender dysphoria diagnosis-mostly based on WPATH Standards of Care, e.7. [39]. The protocol entailed that the diagnostic work up spans at least a six months’ period and incorporates several psychiatric and psychological assessment tools. Various aspects of our clients’ gender identity and sexuality, including their sexual orientation, is scrutinized and the final permission for transition would not be issued pending fulfilment of all the criteria established by international consensus.

Secondly, with regards to the methodological concerns, it is important to note that we “only” performed the present study on a 1.5 Tesla MRI scanner and that 3 Tesla scanners might yield higher SNR. Nevertheless, the present findings serve as an important illustration that subcortical structures or structures close to the edge of the brain may particularly benefit from additional careful steps to increase SNR in any setting. It is also important to note that the test-retest approach to increase the reliability may have its own limitations; staying longer in the scanner and factors such as head motion that increase the noise might be pertinent. Future studies may focus on improvement of MRI sequences or the analysis algorithms and softwares that parse and measure these ROI.

In summary, this study largely replicated findings from prior works in Western cultures and extended them to a Middle Eastern population. Notably, not only the study replicated the important finding of reduced GMV in the right fusiform gyrus in trans men compared to cis women (a finding that was specific to these groups), it also found the right thalamus as an additional region of difference among the two groups. Moreover, using a second scan appeared to improve findings in subcortical structures or regions close to the edge of the brain (frontal poles), thus being relevant for neuroimaging research in other clinical groups with small prevalence rates when anatomical data is of interest [38,39].

## Methods

### Participants

One hundred and forty-one people participated: 40 trans men (TM, who were assigned as female at birth but developed a male gender identity), 41 trans women (TW, who were assigned as male at birth but developed a female gender identity), 30 cisgender women (CW, who were assigned as female at birth and developed a female gender identity) and 30 cisgender men (CM, who were assigned as male at birth and developed a male gender identity). One trans woman had to be excluded given that FreeSurfer software [40,41] was not able to reliably parse the data. One hundred and forty remaining participants were included in the study (full demographics given in Table 1). Transgender participants were recruited through the Transgender Studies Centre at Mashhad University of Medical Sciences (MUMS), Mashhad, Iran. Transgender individuals were interviewed by at least two experienced psychiatrists according to DSM-5 criteria for Gender Dysphoria (GD) (American Psychiatric Association, 2013). Management of GD was based on the Standards of Care (v.7) of the World Professional Association for Transgender Health (WPATH)[42]. Candidate participants were excluded if they had history of head injury (n = 6) or neurological disorder (n =1), used psychotropic medications (n=2), were unwilling to join the study (n = 5) or had history of hormonal therapy (n = 12). Cisgender individuals were recruited through online advertisements in social networks. Anyone interested in participating was then sent an online questionnaire to complete. In this questionnaire certain medical and psychological information were obtained to ensure qualification based on inclusion criteria. Cisgender candidates were excluded if they had history of head injury (n=5) or neurological disorders (n = 3), used psychotropic medications (n=21), had history of hormonal therapy (n=0), did not describe themselves as a cisgender (n=2), or were not heterosexual (n=7). All participants gave a written informed consent to participate in the study. No compensation was made for participation. The study was approved by the ethical committee of Mashhad University of Medical Sciences.

### Psychological questionnaires and interview

In addition to demographic data (e.g., socioeconomic status) and medical histories (surgical, psychiatric and neurological), all participants were asked to score their gender identity and sexual orientation on a scale of 0 to 10. For gender identity, 0 corresponded to feeling fully masculine and 10 to feeling fully feminine. For sexual orientation, 0 represented predominantly attracted to men and 10 predominantly attracted to women (Table 1).

All participants were evaluated regarding their lifetime psychiatric diagnoses. Transgender participants were also interviewed by a psychiatrist using Structured Clinical Interview based on DSM-5 (SCID) [43]. Axis I and axis II disorders were diagnosed if present and reported. Each interview lasted for 90 to 120 minutes.

### MRI acquisition

MRI data were collected on a 1.5T Siemens Magnetom Avanto Syngo MR B17 scanner equipped with an 8-channel phased array receiving coil at Qaem University Hospital of Mashhad University of Medical Sciences. To obtain high quality data the sequence was adapted from the Alzheimer’s Disease Neuroimaging Initiative (ADNI). A T1 weighted MPRAGE sequence was acquired for structural study (160 slices; voxel size: 1.3×1.3×1.2 mm3; FoV: 240 mm; TR: 2400 ms; TE: 3.61 ms; Flip angle: 8°; TI: 1000 ms; acquisition time: 7 min, 42 sec). The adapted ADNI protocol included a MPRAGE Repeat with the same parameters. Both scans were acquired in the same session continuously back-to-back.

### Image preprocessing

Prior to preprocessing, images were inspected for quality checking (NB & HS) and to rule out possible pathologies by a clinician (BA). Data were preprocessed with FreeSurfer (release 5.3.0; Martinos Center for Biomedical Imaging, Charlestown, Mass., USA; http://surfer.nmr.mgh.harvard.edu; [40,41] on the Ghent University High Performance Cluster. The recon-all command was used, which runs the preprocessing steps including bias field correction, intensity normalization, motion correction and skull stripping among others. Then quality control procedures were followed as per ENIGMA protocol (http://enigma.ini.usc.edu/training/) which also involved visual inspection of subject-by-subject outputs to identify possible misclassifications.

## Data analysis

### Neuroimaging data

As noted in the introduction, our ten *a priori* ROIs (bilateral) were selected based on previous literature. This selection sought to balance 1) the groups that have previously published on anatomical differences in transgender persons to avoid implicit biases in region selection, 2) identify regions with inconsistent findings, and 3) used functional data that indicated sensitivity to sex hormones [16,17,26–28] given a hypothesized role of sex hormones in being transgender [44]. However, to keep the number of regions low (to retain sufficient statistical power), a balance between cortical and subcortical regions was sought (Figure 1). A mixed model was fitted for every ROI and a random intercept for each subject. An analysis of variance (ANOVA, with the between subject factor of group: CM, CW, TM, TW) was run using the False Discovery Rate (FDR) to correct for multiple comparisons at the ROI level, i.e., for the number of ROIs (p<.05, FDR corrected)^1^. This was done separately for the independent variables of volume, surface area, and cortical thickness. Regions that showed significant differences on the average scan were further investigated with post-hoc paired t-tests to determine which groups were driving the effects. Bonferroni was then used to correct for multiple comparisons within each ROI. Decision for the respective use of these two methods was made on the notion that FDR seemed more appropriate for the overall number of tests, while Bonferroni seemed best for the post-hoc follow-up. To check whether the analysis incorporating both sets of scans confers gain in SNR compared with an analysis that incorporates only one set, general linear model (GLM) was applied in three separate rounds: once for each set of scans and once in the repeated measures analysis which incorporates both set of scans for each subject and their average; the results where then juxtaposed and compared. Alpha value was set at 0.05 (two-tailed) for all corrected measures.

**Figure 1.**
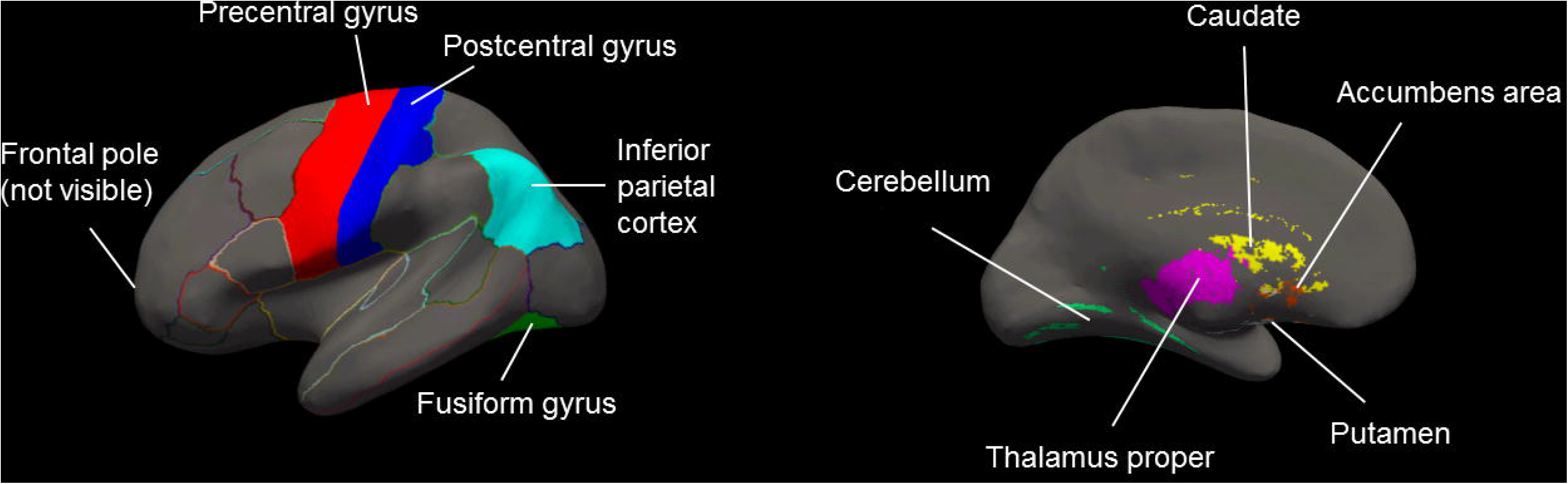
Figure depicts the 10 *a priori* anatomical Regions of Interest (ROIs) balancing cortical and subcortical regions.

**Figure 2.**
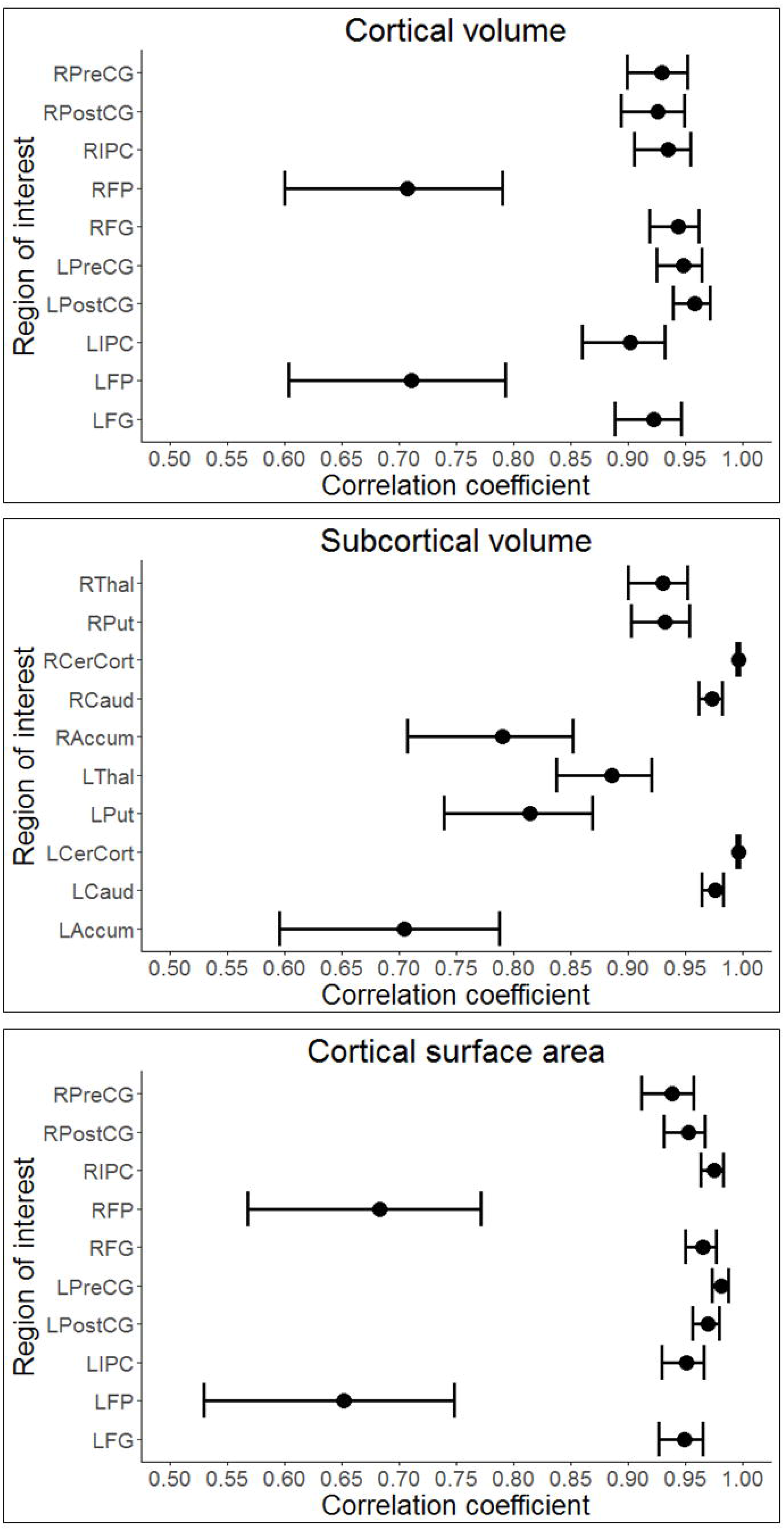
Figure depicts the actual obtained correlation coefficients for each ROI between the two scans for cortical volume (top panel) and subcortical volume (middle panel) as well as cortical surface area (bottom panel). Error bars are 95% confidence intervals. R = right; L = left; PreCG = precentral gyrus; PostCG = postcentral gyrus; IPC = inferior parietal cortex; FP = frontal pole; FG = usiform gyrus; Thal = thalamus; Put = putamen; CerCort = cerebellum; Caud = caudate; Accum = accumbens

None of the 10 ROIs survived the threshold of significance for CTh (with or without using covariates). As such, no post-hoc analysis was performed on this measure. Statistics regarding CTh will therefore no longer be mentioned here (see Table 4 for raw mean/SD).

**Table 4.**
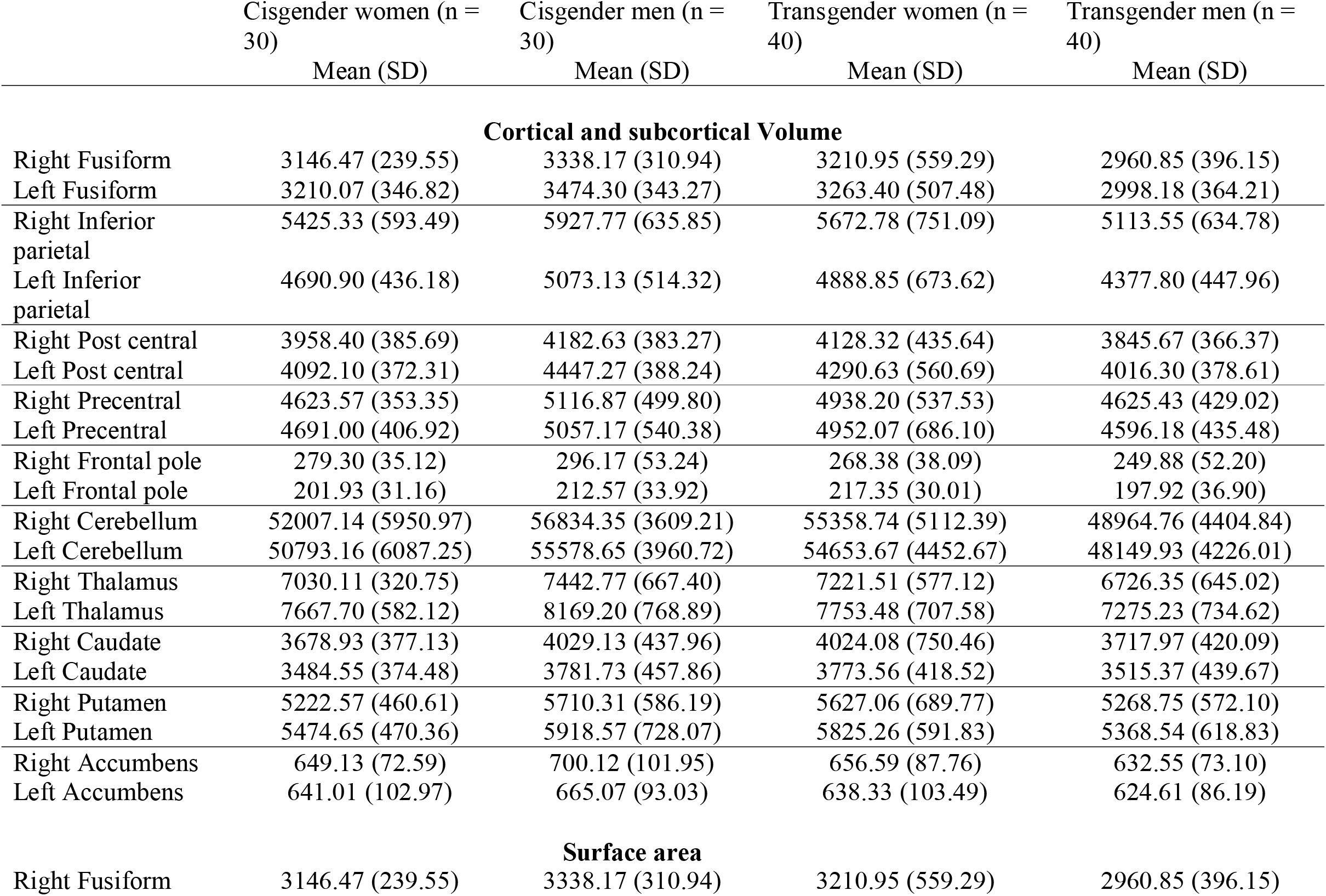
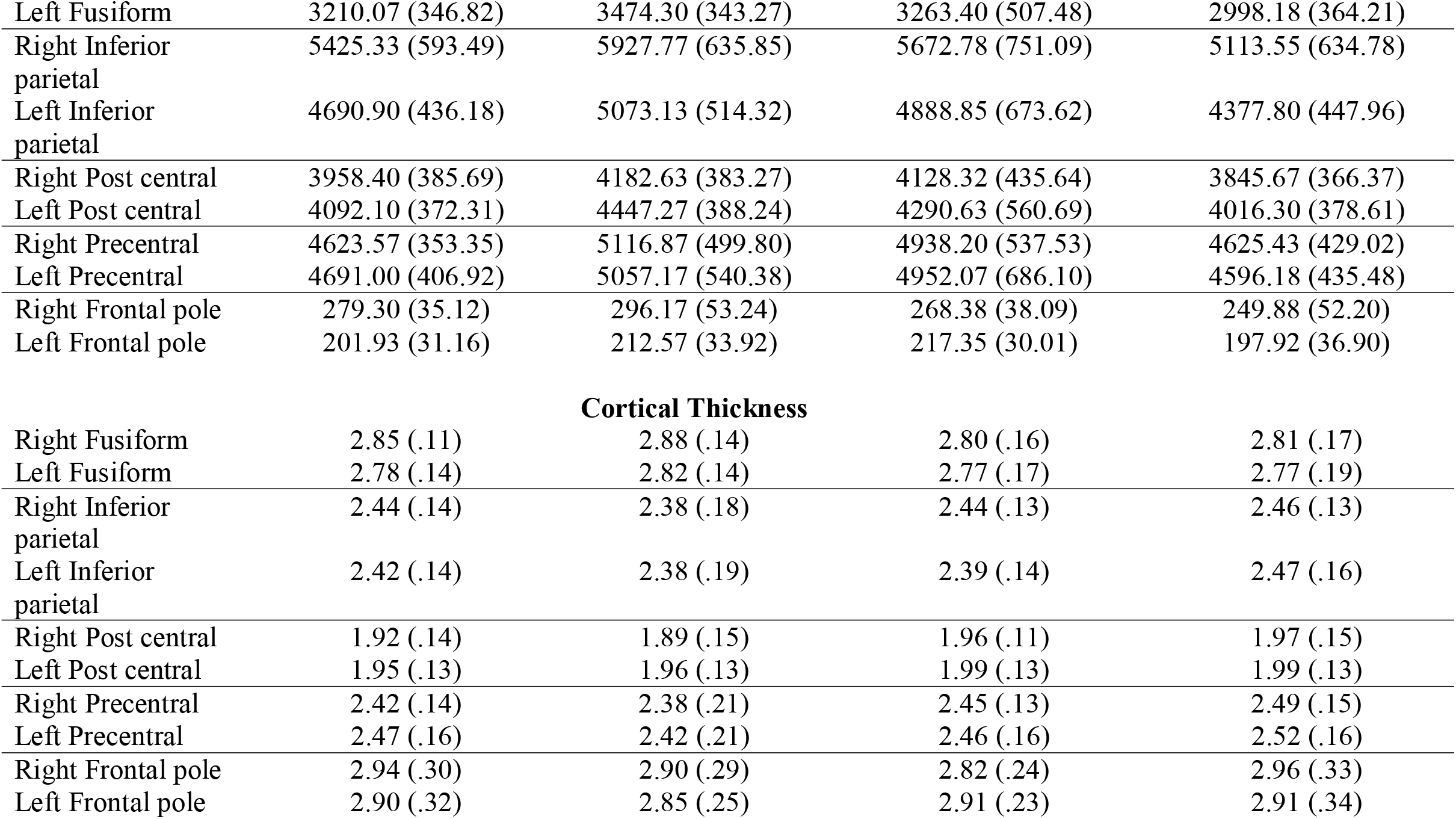
Mean raw extracted values for each of the different measurement types. For volumes, values are in mm^3^, for area values are in mm^2^ and for thickness values are in mm. Please note that for subcortical ROIs no values for ‘cortical thickness’ or ‘surface area’ are available

|  | Cisgender women (n =<br>30) | Cisgender men (n =<br>30) | Transgender women (n =<br>40) | Transgender men (n =<br>40) |
| --- | --- | --- | --- | --- |
|  | Mean (SD) | Mean (SD) | Mean (SD) | Mean (SD) |
| **Cortical and subcortical Volume** |  |  |  |  |
| Right Fusiform | 3146.47 (239.55) | 3338.17 (310.94) | 3210.95 (559.29) | 2960.85 (396.15) |
| Left Fusiform | 3210.07 (346.82) | 3474.30 (343.27) | 3263.40 (507.48) | 2998.18 (364.21) |
| Right Inferior parietal | 5425.33 (593.49) | 5927.77 (635.85) | 5672.78 (751.09) | 5113.55 (634.78) |
| Left Inferior parietal | 4690.90 (436.18) | 5073.13 (514.32) | 4888.85 (673.62) | 4377.80 (447.96) |
| Right Post central | 3958.40 (385.69) | 4182.63 (383.27) | 4128.32 (435.64) | 3845.67 (366.37) |
| Left Post central | 4092.10 (372.31) | 4447.27 (388.24) | 4290.63 (560.69) | 4016.30 (378.61) |
| Right Precentral | 4623.57 (353.35) | 5116.87 (499.80) | 4938.20 (537.53) | 4625.43 (429.02) |
| Left Precentral | 4691.00 (406.92) | 5057.17 (540.38) | 4952.07 (686.10) | 4596.18 (435.48) |
| Right Frontal pole | 279.30 (35.12) | 296.17 (53.24) | 268.38 (38.09) | 249.88 (52.20) |
| Left Frontal pole | 201.93 (31.16) | 212.57 (33.92) | 217.35 (30.01) | 197.92 (36.90) |
| Right Cerebellum | 52007.14 (5950.97) | 56834.35 (3609.21) | 55358.74 (5112.39) | 48964.76 (4404.84) |
| Left Cerebellum | 50793.16 (6087.25) | 55578.65 (3960.72) | 54653.67 (4452.67) | 48149.93 (4226.01) |
| Right Thalamus | 7030.11 (320.75) | 7442.77 (667.40) | 7221.51 (577.12) | 6726.35 (645.02) |
| Left Thalamus | 7667.70 (582.12) | 8169.20 (768.89) | 7753.48 (707.58) | 7275.23 (734.62) |
| Right Caudate | 3678.93 (377.13) | 4029.13 (437.96) | 4024.08 (750.46) | 3717.97 (420.09) |
| Left Caudate | 3484.55 (374.48) | 3781.73 (457.86) | 3773.56 (418.52) | 3515.37 (439.67) |
| Right Putamen | 5222.57 (460.61) | 5710.31 (586.19) | 5627.06 (689.77) | 5268.75 (572.10) |
| Left Putamen | 5474.65 (470.36) | 5918.57 (728.07) | 5825.26 (591.83) | 5368.54 (618.83) |
| Right Accumbens | 649.13 (72.59) | 700.12 (101.95) | 656.59 (87.76) | 632.55 (73.10) |
| Left Accumbens | 641.01 (102.97) | 665.07 (93.03) | 638.33 (103.49) | 624.61 (86.19) |
| **Surface area** |  |  |  |  |
| Right Fusiform | 3146.47 (239.55) | 3338.17 (310.94) | 3210.95 (559.29) | 2960.85 (396.15) |
| Left Fusiform | 3210.07 (346.82) | 3474.30 (343.27) | 3263.40 (507.48) | 2998.18 (364.21) |
| Right Inferior parietal | 5425.33 (593.49) | 5927.77 (635.85) | 5672.78 (751.09) | 5113.55 (634.78) |
| Left Inferior parietal | 4690.90 (436.18) | 5073.13 (514.32) | 4888.85 (673.62) | 4377.80 (447.96) |
| Right Post central | 3958.40 (385.69) | 4182.63 (383.27) | 4128.32 (435.64) | 3845.67 (366.37) |
| Left Post central | 4092.10 (372.31) | 4447.27 (388.24) | 4290.63 (560.69) | 4016.30 (378.61) |
| Right Precentral | 4623.57 (353.35) | 5116.87 (499.80) | 4938.20 (537.53) | 4625.43 (429.02) |
| Left Precentral | 4691.00 (406.92) | 5057.17 (540.38) | 4952.07 (686.10) | 4596.18 (435.48) |
| Right Frontal pole | 279.30 (35.12) | 296.17 (53.24) | 268.38 (38.09) | 249.88 (52.20) |
| Left Frontal pole | 201.93 (31.16) | 212.57 (33.92) | 217.35 (30.01) | 197.92 (36.90) |
| Cortical Thickness |  |  |  |  |
| Right Fusiform | 2.85 (.11) | 2.88 (.14) | 2.80 (.16) | 2.81 (.17) |
| Left Fusiform | 2.78 (.14) | 2.82 (.14) | 2.77 (.17) | 2.77 (.19) |
| Right Inferior parietal | 2.44 (.14) | 2.38 (.18) | 2.44 (.13) | 2.46 (.13) |
| Left Inferior parietal | 2.42 (.14) | 2.38 (.19) | 2.39 (.14) | 2.47 (.16) |
| Right Post central | 1.92 (.14) | 1.89 (.15) | 1.96 (.11) | 1.97 (.15) |
| Left Post central | 1.95 (.13) | 1.96 (.13) | 1.99 (.13) | 1.99 (.13) |
| Right Precentral | 2.42 (.14) | 2.38 (.21) | 2.45 (.13) | 2.49 (.15) |
| Left Precentral | 2.47 (.16) | 2.42 (.21) | 2.46 (.16) | 2.52 (.16) |
| Right Frontal pole | 2.94 (.30) | 2.90 (.29) | 2.82 (.24) | 2.96 (.33) |
| Left Frontal pole | 2.90 (.32) | 2.85 (.25) | 2.91 (.23) | 2.91 (.34) |

## Acknowledgements

The data collection took place in MRI department of Ghaem hospital affiliated to Mashhad University of Medical Sciences. The computational resources (Stevin Supercomputer Infrastructure) and services used in this study were provided by the VSC (Flemish Supercomputer Center), funded by Ghent University, FWO and the Flemish Government – department EWI. The datasets generated and analyzed during the current study are available in the OpenNeuro (https://openneuro.org/) repository, at DOI: 10.18112/openneuro.ds002234.v1.0.0

## Compliance with Ethical Standards

### Funding

This study was funded by the Mashhad University of Medical Sciences.

### Ethical approval

All procedures performed in studies involving human participants were in accordance with the ethical standards of the institutional research committee and with the 1964 Helsinki declaration and its later amendments or comparable ethical standards.

### Informed consent

Informed consent was obtained from all individual participants included in the study.

## Footnotes

1 Previous studies differ on whether they examine absolute or relative volume differences and therefore whether they use some form of correcting for whole brain volume (total brain volume (TBV), total grey matter volume (GMV), intracranial volume (ICV)). One goal of the present study was to compare and replicate the previous findings of Mueller et al. (2017). Since they covaried for age and ICV, the mixed models were re-run with these two covariates included to facilitate direct comparison with that prior study (Supplementary Table 4). Additional analyses with sexual orientation as a covariate did not change the findings.

