## Supplementary Table 1 for "Improving reliability in clinical neuroimaging: a study in transgender persons"

**Supplementary Table 1.** Results for group comparison for grey matter volume (GMV) and cortical surface area (CSA) with age and ICV as covariates included (p<.05, FDR-corrected)

| Main effect of group for GMV | | | | Main effect of group for CSA | | |  |
| --- | --- | --- | --- | --- | --- | --- | --- |
| Region | Scan 1 | Scan 2 | RM- model | Region | Scan 1 | Scan 2 | RM-model |
| L fusiform g. | **<.001** | **.001** | **<.001** | L fusiform g. | .349 | .359 | **<.001** |
| L inf. parietal g. | **.003** | **.001** | **<.001** | R fusiform g. | .514 | .359 | **.001** |
| L postcentral g. | **.025** | **.022** | **<.001** | L inf. parietal g. | .267 | .140 | **<.001** |
| L precentral g. | .230 | .106 | **.008** | R inf. parietal g. | .407 | .359 | **<.001** |
| L frontal pole | **.025** | .116 | **.007** | L postcentral g. | .848 | .632 | **.001** |
| R fusiform g. | **<.001** | **<.001** | **<.001** | R postcentral g. | .909 | .941 | **.012** |
| R inf. parietal g. | **.004** | **.002** | **<.001** | L precentral g. | .621 | .532 | **.001** |
| R postcentral g. | **.009** | .076 | **.007** | R precentral g. | .407 | .359 | **<.001** |
| R precentral g. | **.031** | **.013** | **.001** | L frontal pole | .909 | .814 | .539 |
| R frontal pole | **.002** | **.022** | **.002** | R frontal pole | **.038** | .140 | **<.001** |
| L cerebellum | **<.001** | **<.001** | **<.001** |  |  |  |  |
| R cerebellum | **<.001** | **<.001** | **<.001** |  |  |  |  |
| L thalamus | **<.001** | **<.001** | **<.001** |  |  |  |  |
| L caudate | **.023** | **.044** | **.002** |  |  |  |  |
| L putamen | .070 | .106 | **.004** |  |  |  |  |
| L nucleus accumbens | .569 | .244 | .547 |  |  |  |  |
| R thalamus | **.002** | **.001** | **<.001** |  |  |  |  |
| R caudate | **.002** | **.020** | **.001** |  |  |  |  |
| R putamen | **.001** | **.001** | **<.001** |  |  |  |  |
| R nucleus accumbens | .233 | .269 | .115 |  |  |  |  |

*L: Left, R: Right; inf. =inferior; cereb. = cerebellar; g. = gyrus; RM =repeated-measures; significant p values are in bold font, ICV: Intracranial volume*
