## Supplementary Table 2 for "Improving reliability in clinical neuroimaging: a study in transgender persons"

**Supplementary Table 2.** Two group comparisons for gray matter volume (GMV, in left columns, white background) and cortical surface area (CSA, in right columns, grey background) with age and intracranial volume (ICV) included as covariates. All comparisons p<.05 (Bonferroni corrected).

| Region | CM vs. CW | | CW vs. TM | | TW vs. CW | | CM vs. TM | | CM vs. TW | | TW vs. TM | |
| --- | --- | --- | --- | --- | --- | --- | --- | --- | --- | --- | --- | --- |
| L fusiform g. | .883 | 1 | **.049^b^** | .68 | 1 | 1 | **<.001^d^** | .154 | 1 | .876 | **.001^f^** | 1 |
| L inf. parietal g. | 1 | 1 | .054 | .513 | 1 | 1 | **<.001^d^** | **.016^d^** | 1 | 1 | **.015^f^** | .218 |
| L postcentral g. | 1 | 1 | 1 | 1 | 1 | 1 | **.010^d^** | 1 | 1 | 1 | .233 | 1 |
| L precentral g. | 1 | 1 | 1 | 1 | 1 | 1 | .324 | .666 | 1 | 1 | .184 | .870 |
| L frontal pole | 1 | - | 1 | - | .156 | - | 1 | - | .269 | - | **.019^f^** | - |
| R fusiform g. | .533 | 1 | **.020^b^** | .205 | 1 | 1 | **<.001^d^** | .254 | .718 | 1 | **.008^f^** | 1 |
| R inf. parietal g. | 1 | 1 | .198 | 1 | 1 | 1 | **.005^d^** | **.044^d^** | 1 | 1 | **.002^f^** | .842 |
| R postcentral g. | 1 | 1 | 1 | 1 | 1 | 1 | .322 | 1 | 1 | 1 | **.046^f^** | 1 |
| R precentral g. | 1 | .541 | 1 | .447 | **.047^c^** | .466 | .221 | 1 | 1 | 1 | **.024^f^** | 1 |
| R frontal pole | .381 | .417 | 1 | 1 | 1 | 1 | **.002^d^** | .051 | .286 | **.037^e^** | .701 | 1 |
| L cerebellum | .149 |  | .446 |  | **.012^c^** |  | **<.001^d^** |  | 1 |  | **<.001^f^** |  |
| R cerebellum | .086 |  | .203 |  | .071 |  | **<.001^d^** |  | 1 |  | **<.001^f^** |  |
| L thalamus | 1 |  | .078 |  | 1 |  | **<.001^d^** |  | 1 |  | **.004^f^** |  |
| L caudate | 1 |  | 1 |  | .236 |  | .074 |  | 1 |  | .145 |  |
| L putamen | 1 |  | 1 |  | .588 |  | .693 |  | 1 |  | .157 |  |
| L nucleus accumbens | - |  | - |  | - |  | - |  | - |  | - |  |
| R thalamus | 1 |  | **.032^b^** |  | 1 |  | **.008^d^** |  | 1 |  | **.012^f^** |  |
| R caudate | .870 |  | 1 |  | .474 |  | **.029^d^** |  | 1 |  | .277 |  |
| R putamen | .451 |  | 1 |  | **.005^c^** |  | .065 |  | 1 |  | **.020^f^** |  |
| R nucleus accumbens | - |  | - |  | - |  | - |  | - |  | - |  |

L: Left, R: Right; inf.=inferior; g. =gyrus; - = post-hoc tests were not performed given a lack of main effect; significant p values are in bold font; ^a^CM > CW, ^b^CW > TM, ^c^TW > CW, ^d^CM > TM, ^e^CM > TW, ^f^TW > TM
