## Supplementary Table 3 for "Improving reliability in clinical neuroimaging: a study in transgender persons"

**Supplementary Table 3.** Comparison of p-values for analyses with and without covariates (age, ICV)(significant p-values after FDR correction in bold font). All p-values for cortical thickness were p>.60).

| Region | GMV | | CSA | |
| --- | --- | --- | --- | --- |
|  | Without Covariates | With Covariates | Without  Covariates | With Covariates |
| L fusiform | **<.001** | **<.001** | **.004** | **<.001** |
| L inferior parietal | **<.001** | **<.001** | **.001** | **<.001** |
| L postcentral | **.001** | **<.001** | **.019** | **.001** |
| L precentral | **.012** | **.008** | **.012** | **.001** |
| L frontal pole | **.008** | **.007** | .612 | .539 |
| R fusiform | **<.001** | **<.001** | **.014** | **.001** |
| R inferior parietal | **<.001** | **<.001** | **.003** | **<.001** |
| R postcentral | **.010** | **.007** | .082 | **.012** |
| R precentral | **.001** | **.001** | **.002** | **<.001** |
| R frontal pole | **.002** | **.002** | **.001** | **<.001** |
| L cerebellum | **<.001** | **<.001** |  |  |
| R cerebellum | **<.001** | **<.001** |  |  |
| L thalamus | **<.001** | **<.001** |  |  |
| L caudate | **.003** | **.002** |  |  |
| L putamen | **.006** | **.004** |  |  |
| L nucleus accumbens | .542 | .547 |  |  |
| R thalamus | **<.001** | **<.001** |  |  |
| R caudate | **.002** | **.001** |  |  |
| R putamen | **<.001** | **<.001** |  |  |
| R nucleus accumbens | .121 | .115 |  |  |
